# Guanabenz treatment improves Oculopharyngeal muscular dystrophy phenotype

**DOI:** 10.1101/375758

**Authors:** Alberto Malerba, Fanny Roth, Pradeep Harish, Jamila Dhiab, Ngoc Lu-Nguyen, Ornella Cappellari, Susan Jarmin, Alexandrine Mahoudeau, Victor Ythier, Jeanne Lainé, Elisa Negroni, Emmanuelle Abgueguen, Martine Simonelig, Philippe Guedat, Vincent Mouly, Gillian Butler-Browne, Cécile Voisset, George Dickson, Capucine Trollet

## Abstract

Oculopharyngeal muscular dystrophy (OPMD) is a rare late onset genetic disease affecting most profoundly eyelid and pharyngeal muscles, leading respectively to ptosis and dysphagia, and proximal limb muscles at later stages. A short abnormal (GCG) triplet expansion in the polyA– binding protein nuclear 1 (PABPN1) gene leads to PABPN1-containing aggregates in the muscles of OPMD patients. It is commonly accepted that aggregates themselves, the aggregation process and/or the early oligomeric species of PABPN1 are toxic in OPMD. Decreasing PABPN1 aggregate load in animal models of OPMD ameliorates the muscle phenotype. In order to identify a potential therapeutic molecule that would prevent and reduce aggregates, we tested guanabenz acetate (GA), an FDA-approved antihypertensive drug, in OPMD cells as well as in the A17 OPMD mouse model. We demonstrate that treating mice with GA reduces the size and number of nuclear aggregates, improves muscle force, protects myofibres from the pathology-derived turnover and decreases fibrosis. GA is known to target various cell processes, including the unfolded protein response (UPR), which acts to attenuate endoplasmic reticulum (ER) stress. Here we used a cellular model of OPMD to demonstrate that GA increases both the phosphorylation of the eukaryotic translation initiator factor 2α subunit (eIF2α) and the splicing of *Xbp1*, key components of the UPR. Altogether these data suggest that modulation of protein folding regulation can be beneficial for OPMD and support the further development of guanabenz or its derivatives for treatment of OPMD in humans.

**Significance Statement:** Oculopharyngeal muscular dystrophy (OPMD) is a rare late onset incurable genetic disease characterized by the formation of insoluble aggregates in skeletal muscles. It has been shown that the reduction of aggregates correlates with an improvement of the disease. Here we used a mouse model of OPMD to show that Guanabenz acetate, the active constituent of a marketed but recently discontinued drug for hypertension, decreases the number and the size of aggregates after systemic delivery and improves many aspects of the disease. We also describe experimental evidences explaining the mechanism behind the efficacy of such compound for OPMD.

## Introduction

Oculopharyngeal muscular dystrophy (OPMD) is a late onset rare autosomal dominant muscular dystrophy. It affects 1:100000 people in Europe but it is more common in population groups such as the Bukharan Jews (1/600) and in Quebec (1/1000) likely due to founder effects (1, 2). OPMD is characterized by progressive eyelid drooping (ptosis), swallowing difficulties (dysphagia) and proximal limb weakness. It is caused by a mutation in the *poly(A) binding protein nuclear 1 (PABPN1)* gene that introduces an abnormal expansion of alanine-encoding (GCN)n trinucleotide repeat in the coding region of exon 1 with 11-18 repeats leading to a mutant protein with an expanded number of alanines (expPABPN1, with 11-18 alanines instead of the normal 10) in the N terminal domain (3). PABPN1 is involved in several steps of mRNA and non-coding RNA biogenesis: PABPN1 controls mRNA poly(A) tail length (4, 5) and regulates the use of alternative polyadenylation (APA) sites (6, 7) with a direct effect on mRNA levels and stability. It is involved in splicing regulation (8, 9) and in the long non-coding RNA (lncRNAs) (10) and small nucleolar RNA (snoRNA) processing (11). It also has an important role in poly(A)-mediated RNA decay or export from the nucleus (4) as well as in RNA hyperadenylation (12). Finally PABPN1 has been shown to be required for paraspeckle formation as well as for RNA editing (13).

PABPN1 expression level varies across mouse and human tissues, and is exceptionally low in skeletal muscle (14). Mutated expPABPN1 is prone to form intranuclear aggregates but only in skeletal muscle (15, 16). To date the exact consequence of the presence of these aggregates in muscle still remains poorly understood but recent studies have suggested that expPABPN1 aggregates that also sequesters wild-type PABPN1 in heterozygous patients could itself contribute to phenotypes associated with OPMD by a loss of function mechanism (13, 14, 17–19). This pathological function of PABPN1 aggregates is supported by the fact that higher frequency of aggregates is observed in severe homozygous patients (20). In addition, reducing PABPN1 aggregation by supplementing drugs such as trehalose or other supposed chaperones has consistently demonstrated enhanced cell survival in cell models of OPMD (21–23) and improved muscle weakness in both mouse and *Drosophila* models (2, 24, 25). Apoptosis and deregulation of the ubiquitin-proteasome system have also been proposed as potential downstream pathological events triggered by the aggregates (26, 27).

There is currently no cure for OPMD and one of the only option for patients is the surgical cricopharyngeal myotomy to improve swallowing and surgical treatment of ptosis. Some innovative therapies are under development, including a phase I/IIa cell therapy trial involving autologous myoblast transplantation that has shown improvement in the swallowing capacity of OPMD patients with a cell dose effect (28). Several studies have shown promising results in animal models of the disease, mainly in *Drosophila* and in the A17 mouse model, where expPABPN1 is overexpressed under control of human skeletal muscle actin promoter (HSA1). A gene therapy approach tested in A17 mice showed a strong recovery from the disease (18). As mentioned above, anti-aggregation pharmacological strategies have shown positive results both *in vitro* and *in vivo* in the same A17 mouse model of OPMD (2, 21, 25, 27) and intravenous injection of trehalose is currently being used in a phase II clinical trial (29). Guanabenz acetate (GA), an FDA-approved antihypertensive drug agonist of α2-adrenergic receptors (30), was recently identified to display anti-prion activity through the inhibition of the protein folding activity (PFAR) mediated by the domain V of the large ribosomal RNA, which is a component of the large subunit of the ribosome (31, 32). GA was shown to be effective in the *Drosophila* model of OPMD when provided as oral treatment in the food, where it decreased muscle degeneration and reduced the size of aggregates (24). Deletions of the ribosomal DNA locus (encoding rRNAs mediating PFAR) was also shown to reduce OPMD phenotypes and act synergistically with sub-effective doses of 6-aminophenanthridine, a PFAR inhibitor like GA (24), suggesting a mechanism involving the PFAR (33, 34). Furthermore, GA has been shown to play a beneficial role in models of neurological diseases (e.g. Amyotrophic lateral sclerosis) by acting on the unfolded protein response (UPR). It has been proposed that GA could decrease protein burden in the endoplasmic reticulum (ER) and so increase the efficiency of ER chaperones by prolonging protein translation attenuation during ER stress (35, 36). ER stress condition activates the UPR through different pathways mediated by Grp78 (also called BiP, immunoglobulin-heavy-chain binding protein): first the dissociation of Grp78 from the protein kinase RNA (PRK)-like ER kinase (PERK) activates PERK that phosphorylates the α-subunit of eukaryotic translation initiation factor-2 (eIFα). The phosphorylation of eIFα attenuates the Cap-dependent protein translation and induces the translation of protein stress responses such as activating transcription factor 4 (ATF4) or protein phosphatase 1 regulatory subunit 15A (PPP1R15A). This latter has a crucial role in terminating the UPR pathway by binding to the catalytic subunit of protein phosphatase 1 holoenzyme (PP1c) and dephosphorylating eIF2α. The second branch of the UPR is initiated by the dissociation of Grp78 from inositol requiring protein 1α (IRE-1α), which induces a non-canonical splicing of X-box binding protein 1 *(XBP1)* mRNA producing the active transcription factor XBP1s that in turn triggers the expression of several genes involved in protein folding. The third pathway is operated by the activating transcription factor 6 (ATF6), a receptor that under ER stress and its dissociation to Grp78 is translocated to the Golgi where it is cleaved to produce a fragment which then acts as a transcription factor to promote expression of *XBP1* mRNA and other proteins involved in protein folding. It has been proposed that GA binds to PPP1R15A, thus preventing its binding to PP1c (37, 38) and prolonging eIF2α phosphorylation with the effect of reducing the accumulation of misfolded proteins in the ER (37, 39), but these data are controversial (40, 41).

Here we show that GA supplementation tested in the same dose range used for the marketed hypertensive GA-based drug Wytensin, significantly reduces the number of aggregates, decreases fibrotic collagen deposition and pathological muscle turnover and improves muscle strength in the A17 mouse model of OPMD. As expPABPN1 expression in skeletal muscles induces the expression of ER stress markers, we studied the mechanism underlying the beneficial effects of GA in muscle cells and found that both Ire1α- and PERK-dependent branches of the UPR are affected by GA administration. Altogether, these results identify GA as a positive molecule for OPMD in the mouse model; they also suggest that ER stress plays a role in OPMD and that the modulation of UPR in response to ER stress has therapeutic potential to attenuate PABPN1 aggregate toxicity.

## Results

### Guanabenz effectively reduces PABPN1 aggregates in OPMD muscle cells

GA has been shown to reduce the size of aggregates in a *Drosophila* model of OPMD (24). We first tested GA in the OPMD Ala17 muscle cell model. When differentiated, this myogenic cellular model expresses PABPN1-Ala17, resulting in the formation of a large number of nuclear aggregates (42, 43). GA was added to the cells at day 2 of differentiation and maintained in the medium for 72 additional hours. In these conditions, treatment with 10 μM of GA led to a significant reduction in the number and size of PABPN1-Ala17 aggregates (Fig 1A-C), without altering cell differentiation (**Figure S1A-D**). At lower GA concentrations (ranging from 0.1 to 1 μM), we observed a dose-dependent reduction in the size of PABPN1-Ala17 aggregates but no effect on their number (**Figure S2A-C**).

**Figure 1:**
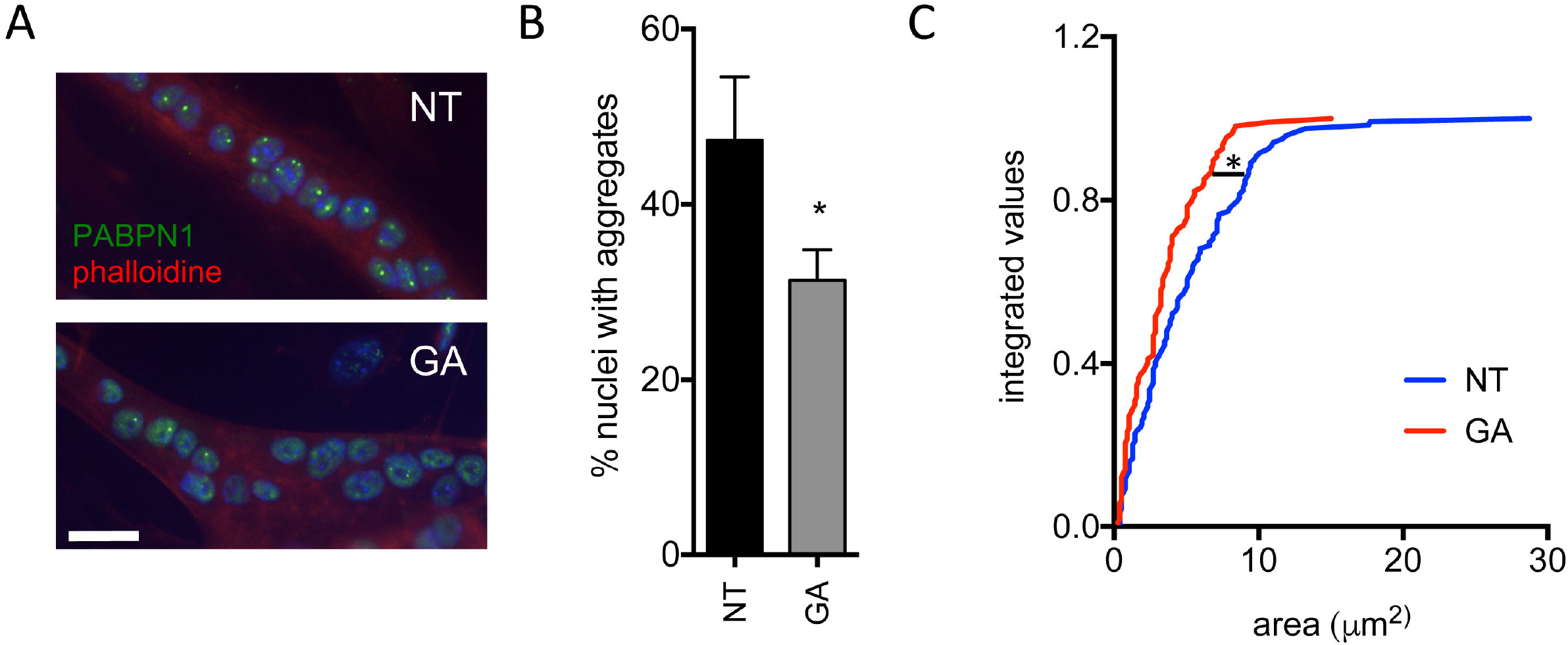
GA treatment reduces PABPN1 aggregates in Ala17 differentiated cells. (**A**) Representative immunostaining of PABPN1 aggregates (green) in differentiated Ala17 cells. Nuclei are counterstained with Hoechst (blue) and myotubes are visualized using phalloidin staining (red). Scale bar = 25 μm. (**B**) The amount of PABPN1 aggregates was reduced upon GA treatment. Unpaired t-test, * p<0.05 n = 3 experiments with at least 100 nuclei counted per experiment. (**C**) The size of PABPN1 aggregates is decreased by GA treatment. Cumulative rank plots show the size of nuclear aggregates in A17 GA-treated group compared to A17 untreated group. Kolmogorov-Smirnov test with *p<0.05. NT= cells treated with DMSO only; GA: cells treated with 10 μM GA.

### ER stress is detected in skeletal muscle of A17 mice

GA is known to affect the UPR in response to cellular ER stress (35–37, 44). No data are available on the ER stress condition and the relative unfolded protein response in muscles of the A17 mouse model of OPMD. Upon ER stress, it has been shown that the lumen of the ER is enlarged (45, 46). Using electron microscopy, we analyzed the skeletal muscle of A17 mice and observed that the muscle ER (sarcoplasmic reticulum, SR) was abnormally expanded in a number of glycolytic fibers suggesting that this cellular organelle is indeed under stress (Figure 2A). In addition, analysis of gene expression profiles of A17 skeletal muscles compared to those of wild-type (WT) mice confirmed that the expression of ER genes (GO term: Endoplasmic reticulum subcompartment) is affected in OPMD mice (Figure 2B). This was further confirmed by qPCR analysis on A17 muscle cDNAs for several crucial UPR-related genes: the chaperones glucose-regulated proteins 78 and 94 *(Grp78, Grp94)*, the DNA damage-inducible transcript 3 *Ddit3* (also named *Chop)*, the activating transcription factor 4 *(Atf4;* downstream of PERK branch) and the DnaJ homolog subfamily C member 3 *(Dnajc3;* downstream of ATF6 branch) (Figure 2C). These data confirm that expPABPN1 expression in skeletal muscles induces the expression of ER stress markers, indicating that the A17 mouse is a relevant OPMD model to evaluate the effect of GA on ER stress and its therapeutic potential *in vivo.*

**Figure 2:**
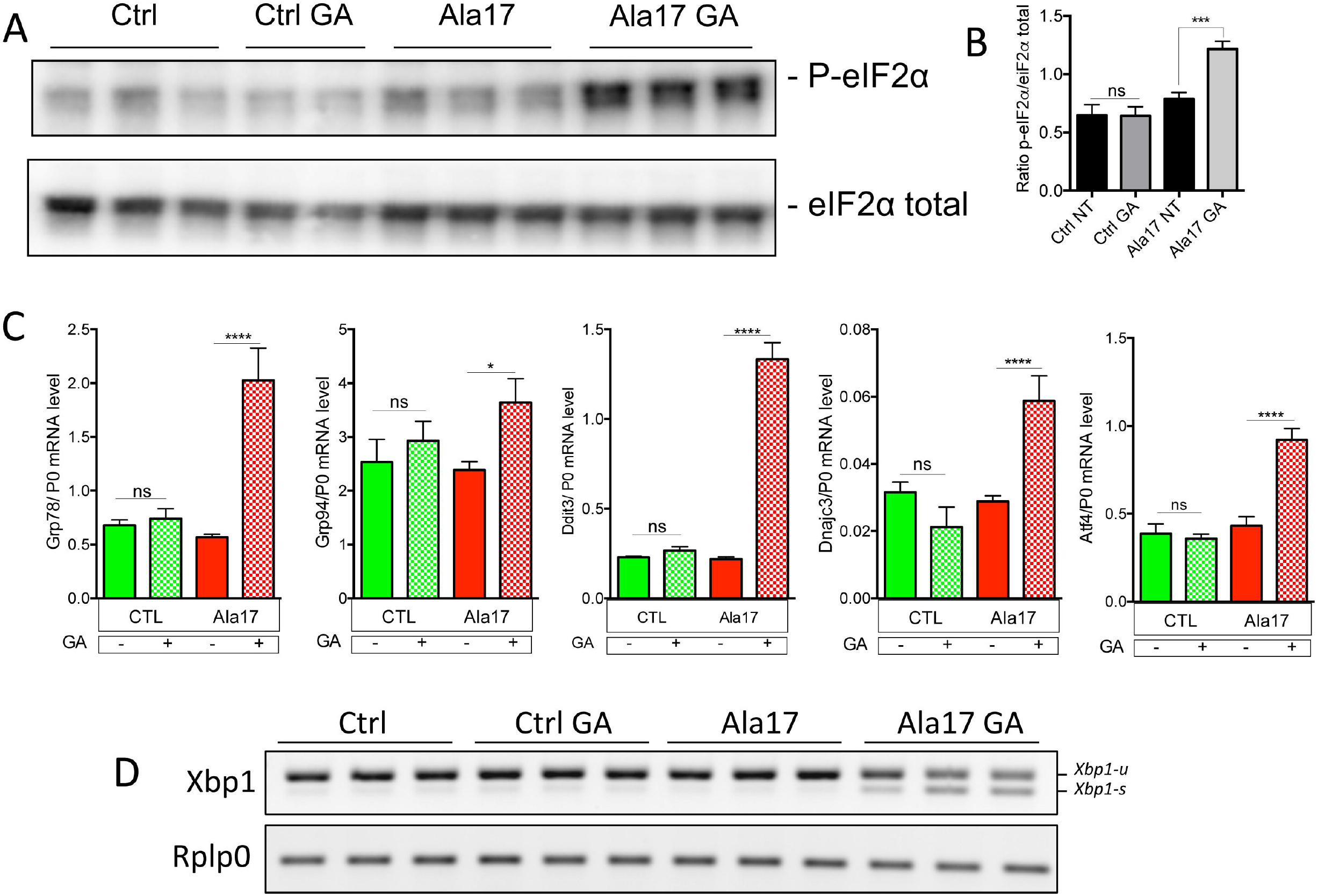
ER stress in the OPMD mouse model. (**A**) Electron micrograph of abnormally enlarged sarcoplasmic reticulum (SR) in A17 mouse. Arrows indicate T-tubules. Scale bar = 2 μm. The boxed area in left panel is enlarged in the right panel. Scale bar = 500 nm. (**B**) Scatter plot of Cellular component ontology on deregulated genes in A17 muscles compared to FvB muscles. The top 30 cellular components are shown in the bubble chart. Gene ratio is shown on the x-axis, point size shows the gene counts and point color shows the adjusted p-value. (C) Key UPR downstream genes *Grp78, Grp94, Ddit3, Dnajc3* and *Atf4* are upregulated in A17 mice compared to control mice. 25 week old mice (n=6 per group) were used for this study. Unpaired t-test **p<0.01, ***p<0.001, ****p<0.0001.

**Figure 3:**
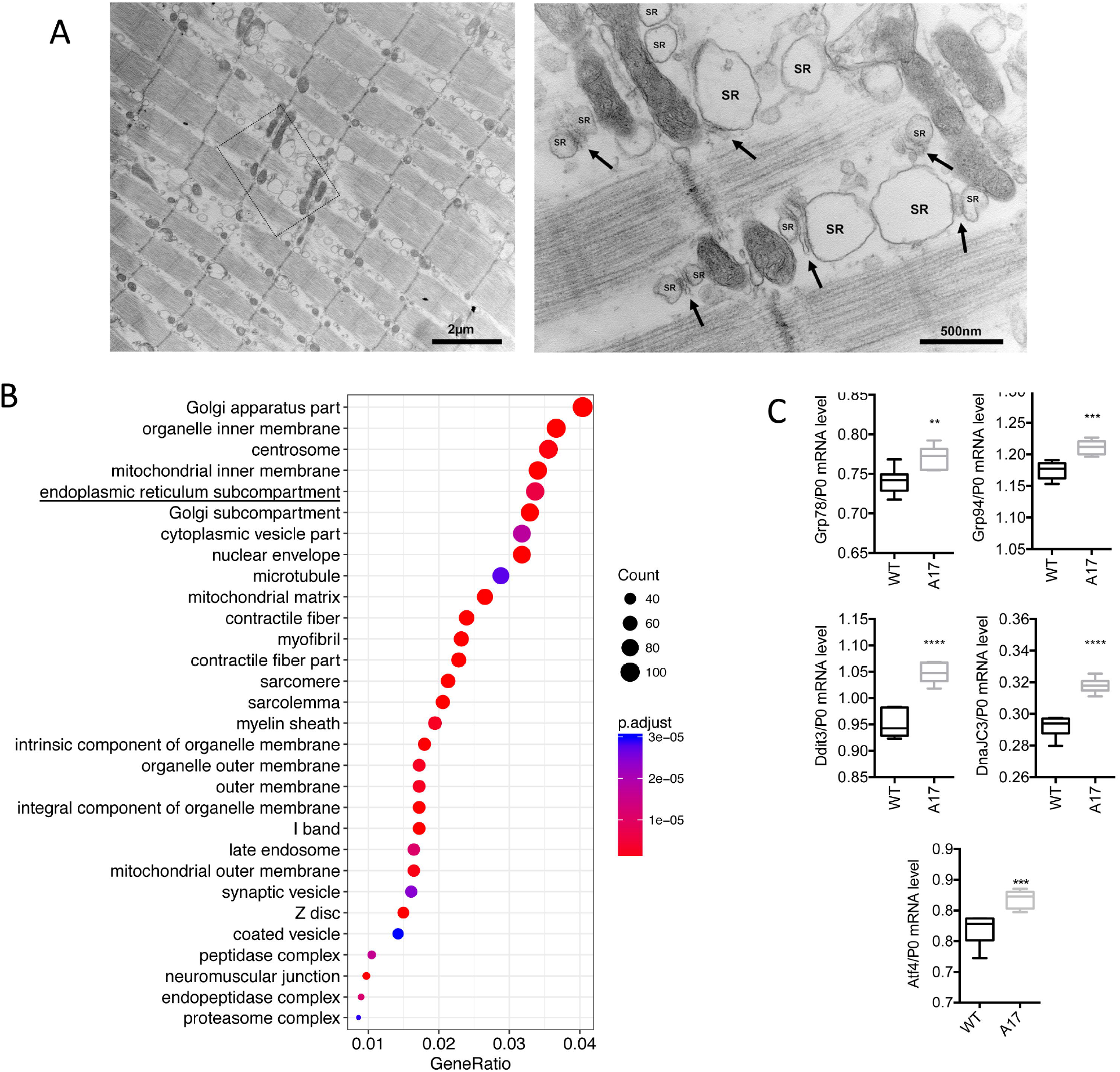
Both PERK-P-eiF2a and IRE1-Xbp1 pathways are activated in response to GA treatment in OPMD cells. (**A**) Western blot showing phosphorylated eiF2α (P-eiF2α) in control and OPMD cells treated with 10 μM GA for 72h (Ala17 GA) compared to DMSO-treated cells (A17). No increase in phosphorylation of eiF2α is observed in control cells treated with GA (Ctrl GA). (**B**) P-eiF2α normalized to total eiF2α. One-way ANOVA test with Bonferroni post-hoc test, ***p <0,001. (C) Downstream key UPR markers are overexpressed in Ala17 cells treated with 10 μM GA (red) while unchanged in control cells (green). mRNA levels were normalized to RPLP0 mRNA level (P0). n=3 per group. One-way ANOVA test *p<0.05, ***p<0.001, ****p< 0.0001 ns: not significant (n=3). (D) 10 μM GA treatment activates *Xbp1* alternative splicing in Ala17 cells but not in control cells, visualized using RT-PCR. The level of total *Xbp1* is comparable in Ala17 cells and in control cells. *Xbp1-u:* unspliced form; *Xbp1* -s: spliced form of *Xbp1.*

### GA treatment induces a favorable change in stress response in Ala17 cells

GA is known to affect the UPR in response to cellular ER stress by prolonging the phosphorylation of the α subunit of eukaryotic translation initiation factor 2 on Ser^51^ (P-eIF2α), a key component of the PERK branch of UPR, to slow down protein translation, decrease the ER protein accumulation in the ER and so leaving more time for the ER chaperones to correctly fold newly synthetized proteins (37). While no effect was observed in GA-treated versus non-treated control cells, a strong increase in phosphorylated eIF2α (P-eIF2α) was observed in GA-treated Ala17 cells (Figure 3A-B) compared to non-treated Ala17 cells. This effect was also observed at lower doses where phosphorylation was activated for all doses ranging from 10 to 0.05 μM of GA (**data not shown**). Significant changes in the expression of the same UPR-related genes were observed in Ala17 treated cells in which *Grp78, Grp94, Ddit3, Atf4* and *Dnajc3* RNA levels were upregulated compared to non-treated cells, while these levels were unchanged in control treated cells (Figure 3C). As Grp78 also activates the IRE1α branch of the UPR, we evaluated if *Xbp1* splicing was affected by GA supplementation. A strong increase in the spliced form of *Xbp1 (Xbp1s)* was detected in GA-treated Ala17 cells (Figure 3D) while no effect was observed on GA-treated control cells. The fact that control cells are not perturbed by GA administration indicates that GA itself does not induce UPR, and is consistent with OPMD cells being challenged by ER stress. Whereas in the animal the expression of the mutated form of PABPN1 induces the UPR by itself, in Ala17 cells GA is needed to reveal UPR activation.

Overall these data show that GA treatment in OPMD cells induces a favorable change in stress response, mediated by increased phosphorylation of eIF2α and Xbp1 splicing activation, which induces UPR-related proteins upregulation. Furthermore, GA treatment only modulates UPR in cells affected by ER stress.

### Treatment with GA in the A17 OPMD mouse model efficiently improves muscle strength, decreases pathological aggregates accumulation, myofibre turn-over, and fibrosis

To test the effect of GA administration in the OPMD mouse model, 10-12 week old A17 mice were injected intraperitoneally (i.p.) with 4 mg/kg of GA (n=6) or with saline (n=6) three times per week (for a final dose of 12 mg/kg/week of GA) for 13 weeks. We chose this dose because it is in the range of the dose of drug used for the marketed GA-based Wytensin to treat hypertension in humans (47). Age-matched WT mice (FvB) (n=6) were also injected with saline as controls. At the end of the experiment, all mice gained a similar bodyweights irrespective of the treatment (**data not shown**). As GA has been shown to induce dizziness, weakness and sedation as possible side-effects (48), locomotor behavior of mice was examined by open field behavioral monitoring both after the first and before the last drug administration in the 13-week long study. After the first injection of GA (or saline in control mice), behavior of mice was monitored for 60 minutes and several parameters describing the animal activity were recorded. This analysis showed that mice treated with GA were less active than mice injected with saline soon after GA administration (One-way ANOVA test, **Figure S3A**). However, the same analysis performed in mice just before the last injection of the study (with mice that were not under the effect of the drug at the time of the analysis) indicated that animals chronically treated with GA for 13 weeks are as active as mice injected with saline (One-way ANOVA test **Figure S3B**) suggesting no detrimental effect of a prolonged treatment with GA in locomotor activity of these mice. Before harvesting the samples from the treated mice, the forelimb force was analyzed by grip strength analysis. A17 mice have weaker forelimbs strength than FvB WT mice (25, 49). The treatment with GA significantly increased the muscle strength of treated A17 mice compared to saline injected A17 mice (Figure 4A). To assess more precisely the muscle strength, TA muscles of saline and GA-treated mice were then analyzed by *in situ* electrophysiology force measurement. Muscles were stimulated with 9 consecutive trains of impulses from 10 to 180 Hz and the muscle strength provided after each train of stimulation was measured. GA treatment significantly increased the absolute maximal tetanic force generated by the muscles of A17 mice (Figure 4B-C). Normalization of the maximal tetanic force with the cross-sectional area of the muscles provides a measure called maximal specific force, which is an indication of the muscle strength per unit of the muscle. Administration of GA to A17 mice significantly increased the maximal specific force and normalized the muscle strength to the WT level (Figure 4D). No difference in muscle weight was observed for tibialis anterior (TA), extensor digitorum longus (EDL), soleus (Sol) or gastrocnemius (Gast) (**Figure S4**). As GA is expected to reduce the amount of aggregates, muscles were analyzed for the presence of PABPN1 aggregates. The treatment with GA significantly reduced both the percentage of nuclei containing aggregates (one-way ANOVA test, from ~31% to ~22% of nuclei) and the size of aggregates in TA skeletal muscles of A17 mice (Figure 5A-C), while the total level of PABPN1 protein was unchanged (Figure 5D-E). Interestingly the proportion of centrally nucleated myofibres, which is an index of muscle degeneration and regeneration, was reduced by almost 50% in GA treated TA muscle (from 7.5% to 4%, Figure 6A) suggesting a reduction on the pathology-derived myofibre turnover. As the OPMD pathology is characterized both in humans and in the A17 mouse model by increased fibrosis and collagen deposition in affected muscles, we stained the FvB and A17 muscles for markers of fibrosis (ie Collagen I, III via Sirius red staining, and Collagen VI via immunostaining). As expected, saline-treated A17 muscles were more fibrotic than age matched saline treated FvB muscles. GA treatment significantly decreased the amounts of collagen proteins detected compared to the wild type levels (Figure 6B-C). Overall these data show that GA administration is beneficial in OPMD muscles where it improves muscle strength, and reduces aggregates formation, cell turn-over and fibrosis with no significant side-effects.

**Figure 4:**
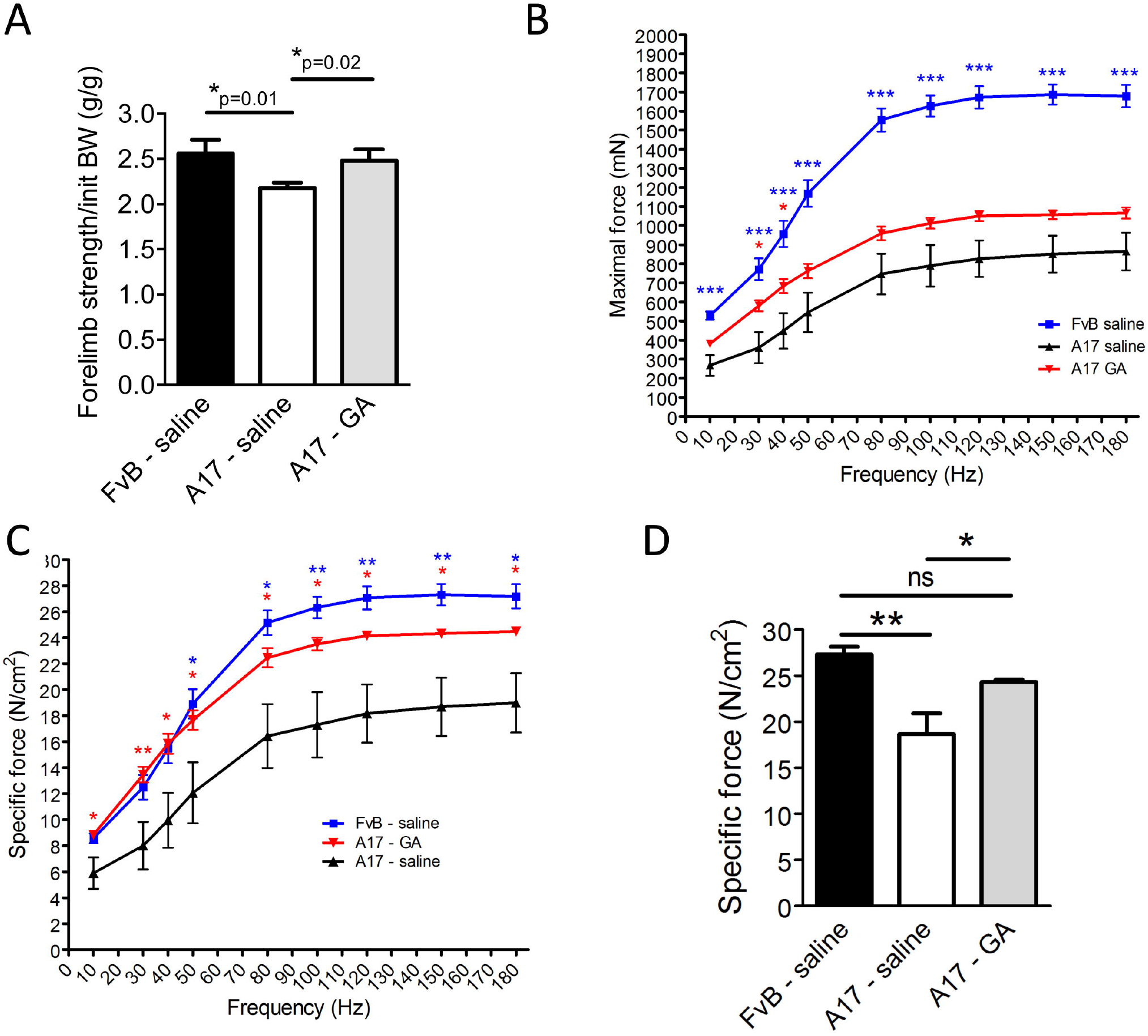
in vivo administration of GA improves the functionality of treated muscles. (**A**) Grip-strength analysis performed in mice at the end of the 13-week long treatment shows that forelimbs of mice treated with GA are significantly stronger than those of mice treated with saline. Unpaired t-test, *p<0.05. (**B**) *in situ* muscle physiology in muscles: force-frequency analysis of maximal force generated by TA muscles of treated mice shows that GA administration significantly increases the maximal force generated by TA muscles. One-way Anova test with Bonferroni post-doc test, *p<0.05, ***p<0.001. (**C**) Normalization of maximal force by muscle cross sectional area provides a measure of the muscle strength per unit of skeletal muscle called specific maximal force. One-way ANOVA test with Bonferroni post-doc test, *p<0.05, **p<0.01. (**D**) Specific maximal force of muscles treated with GA was normalized to the level of control wild type muscles (150 hz stimulation is shown). Mean ± s.e.m., n=6; Oneway ANOVA test with Bonferroni post-doc test, *p<0.05, **p<0.01, ns: not significant. Red and blue stars refer to A17 GA vs A17 saline or FvB saline vs A17 saline comparisons respectively.

**Figure 5:**
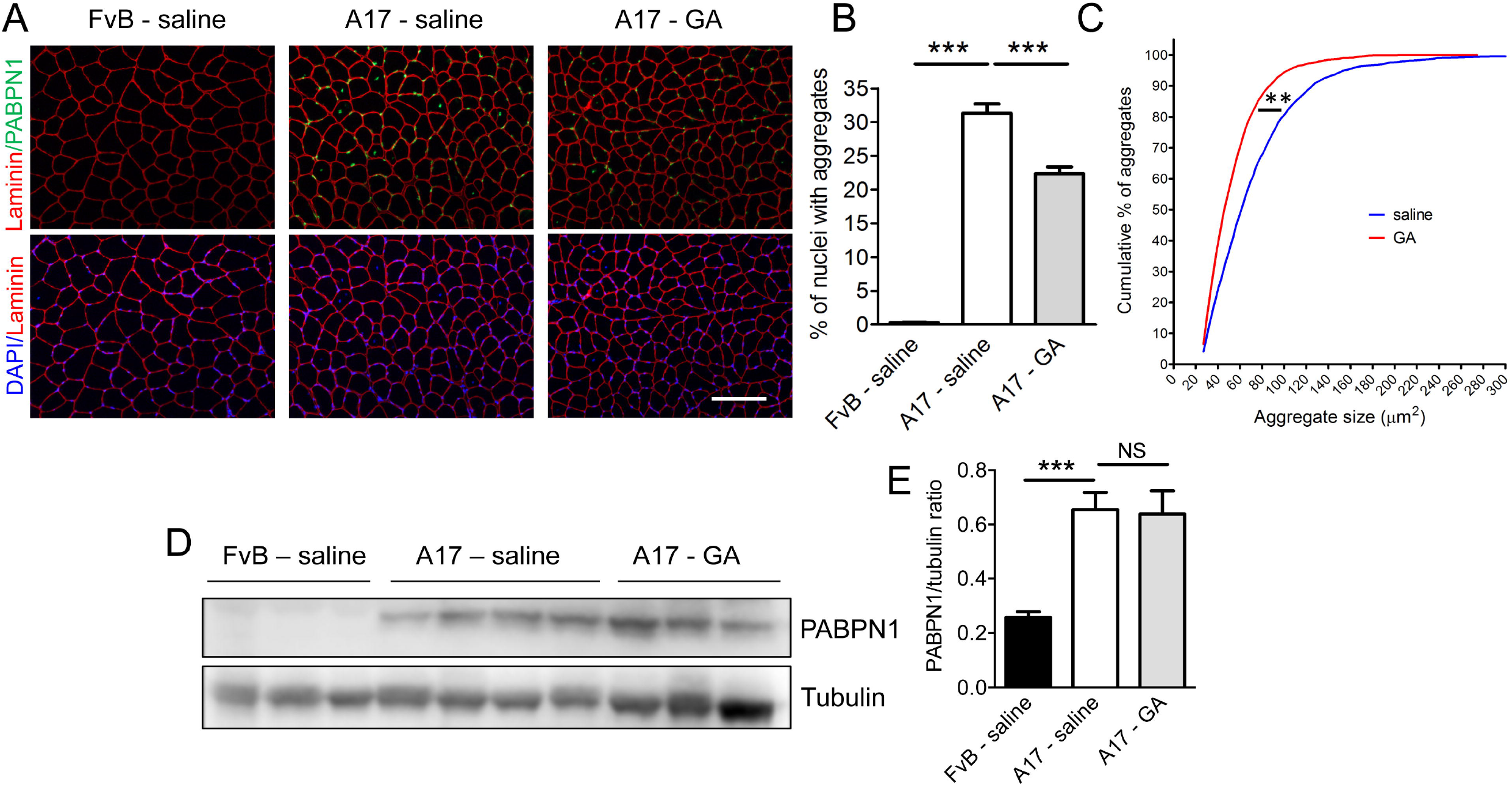
Intraperitoneal injection of GA reduces the amount and the size of intranuclear inclusions. After thirteen weeks of treatment by i.p injection of GA, TA muscles of treated mice were collected and PABPN1 expression was analyzed. (**A**) Detection of PABPN1 intranuclear aggregates (green) and laminin (red) by immunofluorescence in sections of treated muscles. Sections were pre-treated with 1 M KCl to discard soluble PABPN1 from the tissue. Nuclei are counterstained with DAPI (blue). Scale bar = 200 μm. (**B**) Quantification of nuclei containing aggregates in muscle sections indicates that treatment with GA significantly reduces the amount of aggregates to ~20% compared to A17 untreated mice (~35% nuclei containing aggregates). One-way ANOVA test with Bonferroni post-hoc test ***p<0.001. (**C**) GA administration reduces the size of intranuclear aggregates. Cumulative rank plots show the size of aggregates in A17 GA-treated group compared to A17 untreated group. Kolmogorov-Smirnov test with ** p<0.01. (D) Western blot for PABPN1 expression shows that GA treatment does not change the level of PABPN1. (E) Densitometric analysis of western blot for PABPN1 detection shows no difference in PABPN1 expression in saline and GA-treated A17 mice normalized to the relative tubulin expression. n=6 in each group. Data are presented as mean ± s.e.m. One-way ANOVA test with Bonferroni post-hoc test ***p<0.001, ns, not significant.

**Figure 6:**
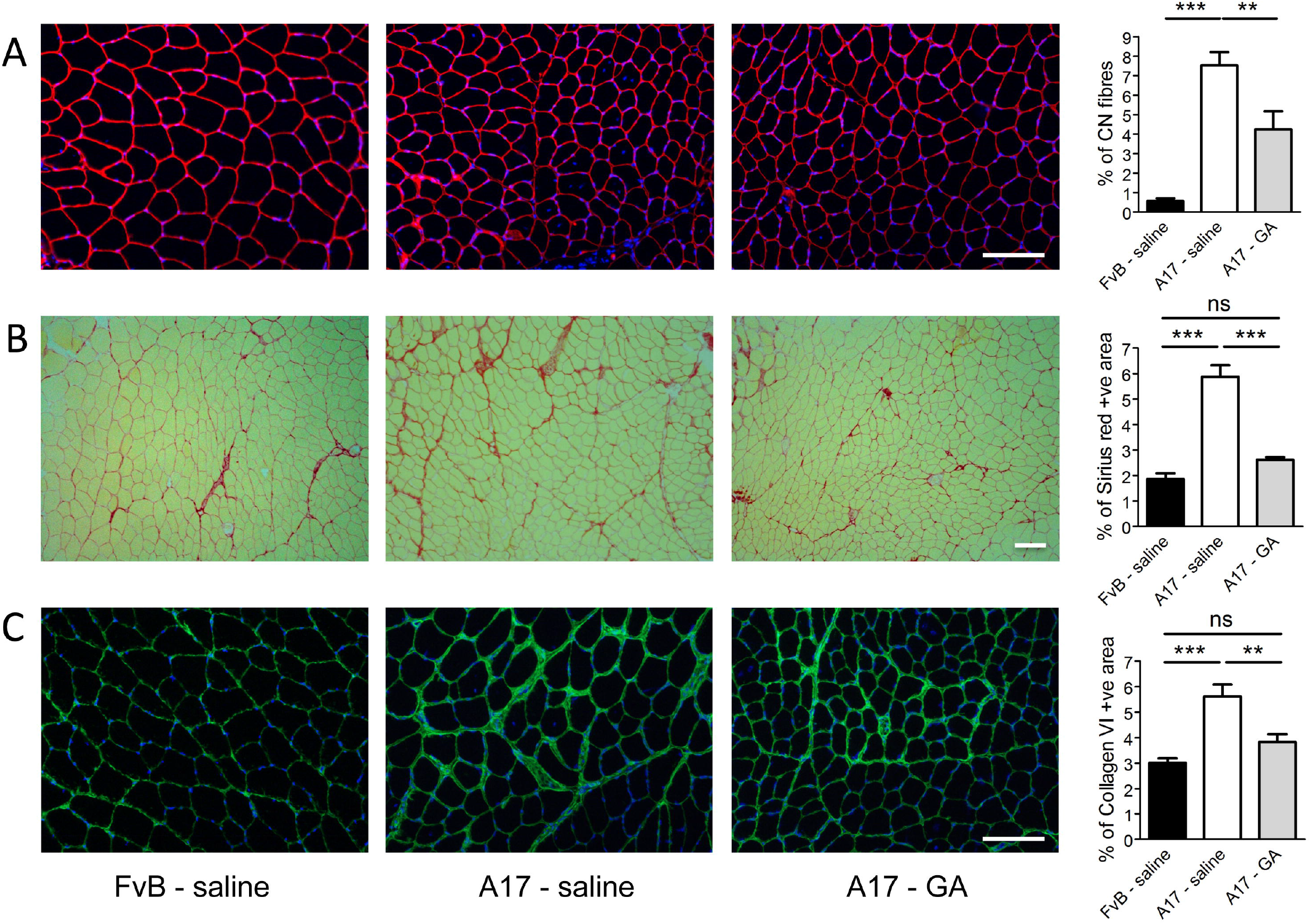
GA administration reduces the percentage of centrally nucleated fibres and the fibrosis in muscle. (**A**) Representative images of histological stainings for Laminin and DAPI were used to quantify the percentage of myofibres with central nuclei. GA treatment significantly reduces the percentage of centrally nucleated fibres (CN) compared to saline treated A17 mice (right panel). One-way ANOVA test with Bonferroni post-hoc test **p<0.01, ***p<0.001. (**B**) Representative images of the Sirius red staining used to quantify the area covered by Collagen I and III in TA muscles of treated mice. Muscle of mice treated with GA have significantly less Collagen I and III positive area compared to muscles of mice treated with saline (right panel). One-way ANOVA test with Bonferroni post-hoc test, ***p<0.001, ns, not significant. (**C**) Representative pictures of Collagen VI immunostaining used to quantify the collagen VI positive (+ve) area. Collagen VI +ve area is smaller in GA-treated muscles compared to saline-treated muscles (right panel) One-way ANOVA test with Bonferroni post-hoc test **p<0.01, ***p<0.001, ns, not significant. Scale bar = 200 μm. n=6 in each group. Data are presented as mean ± s.e.m.

### Oral administration of GA increases muscle strength and reduces the amount of nuclear aggregates of OPMD mice

Since the GA-based drug Wytensin has been authorized for oral prescription in tablet form, we tested the effect of GA administered orally in OPMD mice. GA was tested in 10-12 week old A17 mice (n=6) by formulating the drugs in food pellets to provide a final dose of 12 mg/kg of GA per week. Groups of A17 and FvB male mice (n=7 and 8 respectively) were also fed with standard chow pellets as control. Food was provided *ad libitum*, weighed and changed twice a week. During the experiment, mice fed with GA-supplemented chow ate 15% less food than mice provided with standard chow possibly due to the unpleasant taste of the drugs. However, at the end of the experiment all mice had gained a similar amount of weight (**Figure S5A-B**). As performed for the intraperitoneally-delivered drug experiments, we assessed whether GA affects locomotor activity of the mice and again no difference was observed between control groups and GA-treated mice for any of the parameters analyzed at the end of the treatment period (**Figure S6**). Analysis of PABPN1^-^ positive aggregates showed a decrease in both the percentage of nuclei containing aggregates and the size of the aggregates in muscles treated with GA compared with muscles of control mice (Figure 7A-C). However we did not observe a change in the percentage of centrally nucleated fibres (data not shown). As for the experiment based on i.p. administration, no change was observed in the muscle weight of TA, EDL, Sol and Gast at the end of the study (**data not shown**). Forelimb strength showed a strong trend towards an improvement in A17 mice treated with GA compared with control mice (unpaired t-test: *p=0.038) and was essentially normalized to the level of WT FvB mice (Figure 7D). Overall these results indicate that GA provided orally is also effective in A17 mice by increasing the muscle strength and significantly reducing the size and number of aggregates.

**Figure 7.**
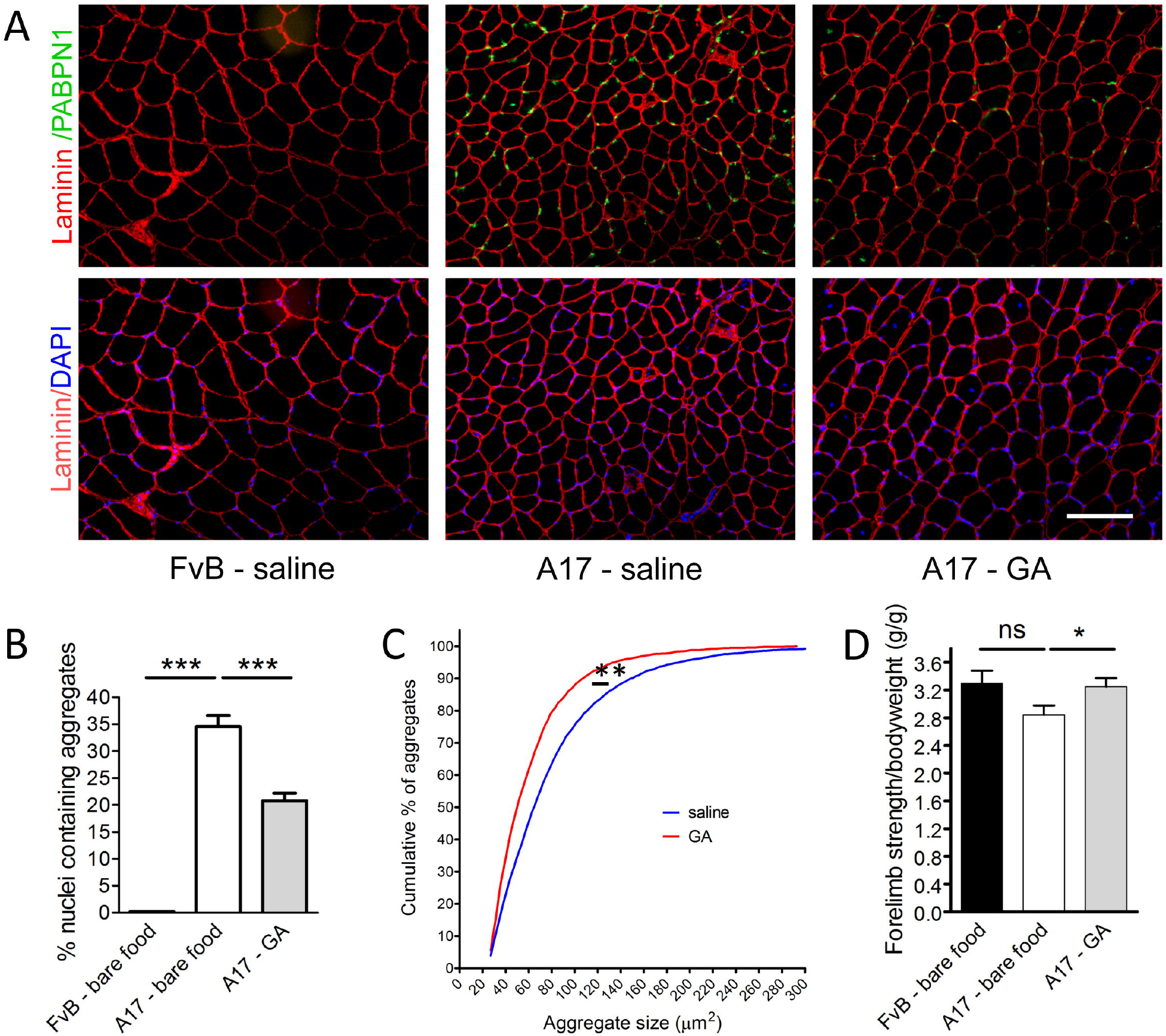
GA formulated in food has beneficial effects in forelimb strength and reduces the amount and the size of intranuclear aggregates. (**A**) Detection of PABPN1 nuclear aggregates (green) and laminin (red) by immunofluorescence in sections of treated muscles. Sections were pre-treated with 1 M KCl to discard all soluble PABPN1 from the tissue. Nuclei are counterstained with DAPI (blue). Scale bar = 200 μm. (**B**) Percentage of nuclei containing aggregates in muscle sections indicates that the treatment with GA significantly reduces the amount of aggregates to ~20% compared to A17 untreated mice. One-way ANOVA test with Bonferroni post-hoc test ***p<0.001 (**C**) GA administration reduces the size of intranuclear aggregates. n=6 per group. Cumulative rank plots show the size of aggregates in A17 GA-treated group compared to A17 untreated group. Kolmogorov-Smirnov test with ** p<0.01. (**D**) Forelimb force of A17 mice treated with GA showed a trend towards an increase in strength compared to the force of mice treated with saline. Unpaired t-test, *p<0.05. ns, not significant. Data are presented as mean ± s.e.m. n=6-8 per group.

## Discussion

OPMD is a late onset disease associated with the formation of intracellular aggregates forming nuclear inclusions in affected tissues, similar to what is found in other adult neurological proteinopathies like Huntington’s, Parkinson’s, Alzheimer’s disease, frontotemporal dementia, spinocerebellar ataxias, inclusion body myositis, spinal and bulbar muscular atrophy, myotonic dystrophy and prion diseases like Creutzfeldt-Jakob (50–52). At present, the contribution of PABPN1 aggregates to the OPMD phenotype is still a subject of debate, but it is commonly accepted that aggregates, the aggregation process and/or the early oligomeric species of expPABPN1 are toxic and crucially involved in the OPMD pathology (2). The development of pharmacological molecules targeting these nuclear aggregates may therefore provide a valid approach for the treatment of OPMD. Several drugs have demonstrated positive effects in preclinical models of OPMD, such as 6-aminophenanthridine (6AP) and GA in *Drososphila* (24), and doxycyclin (25), cystamine (27) and trehalose in mice, with the latter progressed to clinical trial stages in OPMD patients (NCT02015481 on clinicaltrial.gov). In this study, we tested the effect of GA in a cellular model of OPMD and its therapeutic potential in a relevant mouse model of the disease. First of all, we showed that GA decreases the percentage and size of nuclear aggregates both *in vitro* and *in vivo.* GA and 6AP have previously been shown to reduce the size of aggregates and improve the pathology in a *Drosophila* model of OPMD (24). However, while 6AP was shown to use the ribosome-borne protein folding activity of the large ribosomal RNA (PFAR) mechanism to reduce the size of aggregates, the mechanism of action of GA has not been addressed. Several reports, in the context of other diseases, have shown that GA promotes the UPR to counteract ER stress (35, 36, 39). It has been proposed that GA binds the regulatory subunit of protein phosphatase 1 (PPP1R15A), thereby preventing the dephosphorylation of eiF2α and then protein translation recovery following stress (37, 39), although its exact mechanism of action remains controversial (41, 53). Delaying protein translation recovery may thus allow the UPR downstream chaperones (e.g. GRP78, GRP94) to have more time to cope with the impaired ER homeostasis. Here we show that GA promotes UPR in models of OPMD by acting through the UPR pathway initiated by PERK. In response to GA supplementation, we observed both an increase in phosphorylation of eiF2α and an increase in *Xbp1* splicing. It has been described that *Xbp1* splicing is regulated by activation of both PERK (through increased expression of Atf4 that is known to enhance *Xbp1* splicing (54)) and IRE1α pathways. A cross talk between PERK and IRE1α has been previously demonstrated for miR-30c-2-3p expression that is activated by PERK/NF-ĸB and inhibits the IRE1-dependant transcription factor *Xbp1* (55, 56). Here, instead, we show that GA supplementation *in vitro* increases expression of Atf4 (mediated by PERK pathway) that is known to enhance *Xbp1* splicing (54). It has been shown that GA on its own is not able to modulate UPR related genes in unstressed cells (37). We found that cells expressing expPABPN1 did not show a different expression of the markers we studied to assess UPR activation compared to control cells. However, Ala17 cells clearly responded to the treatment with GA by upregulating such markers, suggesting that pre-existing ER stress is present in OPMD cells and that this allows a response to GA treatment. A possible explanation for the lack of detection of these UPR signatures in untreated Ala17 cells is that the expression of expPABPN1 in OPMD myotubes is not long enough to allow detection of UPR response. This is further supported by the observation that in the A17 OPMD mouse model, where expPABPN1 is expressed for a longer time, UPR activation is readily detectable. Although our study strongly suggests a connection, the link between PABPN1 aggregates and ER stress in OPMD remains undefined. OPMD is characterized by mitochondrial, ubiquitin proteasome system (UPS) and apoptosis defects (26, 49, 57, 58), all of which can potentially generate ER stress. Furthermore, protein misfolding in the ER, nutrient deprivation, oxidative stress, calcium disturbance, shortage of chaperone proteins and ageing can also trigger ER stress (Integrated stress response, ISR) (59–63) and all these features are likely present in OPMD cells. For example it is possible that unresolved ER stress generates more substrates for the UPS that would consequently lead to the overwhelming and saturation of the UPS degradation machinery in a self-reinforcing pathological cycle (64). Alternatively trapping protein chaperones like HSP70 in the aggregates (65, 66) would make them less available for normal cellular functions, which could lead to ER stress. Recently, it has been demonstrated that PABPN1 protein regulates nuclear structure and homeostasis of Ca^2^+ (67). The ER is an essential intracellular organelle for many cell functions including the maintenance of Ca^2+^ homeostasis. Calcium is one of the key regulators of cell survival and can induce ER stress-mediated apoptosis (68, 69) mainly through mitochondrial cell death (70). It has been shown that PABPN1 mutation in OPMD gives rise to multiple Ca^2^+ defects such as alterations of the sarcoplasmic reticulum (SR) Ca^2^+ content, inhibition of the voltage-gated SR calcium release and reduced expression of ryanodine receptor (67). Moreover, the presence of *TNNT3* pre-mRNA in nuclear aggregates (42) is associated with aberrant calcium sensitivity in muscle fibres. These studies show that disturbance of Ca^2+^ homeostasis plays a role in OPMD physiopathology and suggests that this could be one of the major causes of ER stress in OPMD. Furthermore OPMD has been proposed as a premature-ageing disease (17) and ER stress is prone to be activated in aging since age-related changes such as increased oxidative stress, protein misfolding, impairment of calcium homeostasis and disturbance of protein synthesis are observed when the protein quality is altered (71). Similarly, the enrichment analysis of transcriptomic data from the OPMD mouse revealed ER as one of the top-five deregulated cellular components.

Following the positive effect of GA described in the *Drosophila* model of OPMD, this study is the first demonstration of the effectiveness of GA in a mammalian model of OPMD. Here, we decided to dose the mice with 12 mg/kg/week as this translates, after conversion for the body surface area, to approximately 10 mg/day in a 70 kg human being which is in the mid-range for the administration of Wytensin, a recently discontinued anti-hypertensive drug whose active ingredient is GA (47). GA i.p. treatment efficiently reduced the percentage of nuclei containing aggregates while simultaneously decreasing muscle fibrosis and improving muscle strength. Furthermore by providing the drug via oral feeding, we demonstrated that the drug was still active and able to decrease the percentage of aggregates using the oral route of delivery, which is compatible with clinical application in humans. A direct comparison with trehalose, the only anti-aggregate therapeutic strategy that has yet progressed to clinical application for OPMD, is impossible due to the different doses, route of delivery and outcome measures used to study these compounds. However the efficacy on reducing the amount of aggregates and the functional muscle improvement seem to be at least comparable if not substantially better after administration of GA, and with the further advantage that GA is already a well-characterized drug that entered the market although for a different pathological condition. This drug (ie Wytensin) was discontinued mainly due to some side-effects (e.g. dizziness and lethargy) that we also observed in the current study soon after dosing the mice. However, our study demonstrates that the chronic administration with GA, at least at the doses used, does not affect the locomotor activity in the long term. In conclusion, we suggest that GA ameliorates OPMD acting, at least partially, by finely modulating the ER stress response and reducing the aggregate formation. GA and novel GA derivatives thus have significant therapeutic potential for translation into clinical trials for the treatment of OPMD.

## Materials and methods

### Drug preparation

For *in vitro* experiments, Guanabenz acetate (GA) (G-110, Sigma) was prepared in DMSO at a stock concentration of 100 mM. Fresh aliquots were used for each new experiment at concentration ranging from 0.1 to 20 μM. For the intraperitoneus (IP) injection, GA was first prepared in 5% DMSO and then diluted to the final concentration at 4 mg/kg of bodyweight with saline (NaCl 0.9% in water). For the oral feeding experiments, food supplemented with GA was prepared using the 2016.12 based diet provided by Envigo.

### Cell culture

Primary immortalized mouse myoblasts H2KB IM2 (control cells) are derived from conditionally immortalized mice using a temperature-sensitive SV40 large T-antigen (tsA58) transgene (72). Mutated PABPN1 (Ala17) and wild-type PABPN1 (Ala10) stable cell lines have been established from IM2 cells transfected with a plasmid containing either mutated or wild type PABPN1 under a desmin promoter driving myotube-specific expression (43). Cells were cultivated on matrigel coated surface in DMEM supplemented with 20% of fœtal calf serum (Invitrogen), 0.5% of chick embryo extract (Seralab), 100 U/ml penicillin-streptomycin antibiotic (Thermofisher) and 10 U/ml interferon-gamma (Millipore) at 33°C in a humidified 5% CO2 air atmosphere. For Ala10 and Ala17 cell lines containing a resistance gene, 500 μg/ml de geneticin (G418; Life Technologies) was added to the media. At 90% of confluency, cells were differentiated in DMEM supplemented with 10% horse serum and 1% penicillin-streptomycin at 37°C in a 5% CO_2_ humidified atmosphere.

### PABPN1 immunostaining in cells

For immunostaining, H2KB cells were cultivated in Ibidi 35 mm Dishes (Biovalley). Immunostaining of PABPN1 was performed at 5 days of differentiation. Cells were washed once in PBS and fixed for 10 min with paraformaldehyde-PBS. Cells were then incubated for 15 min in PBS-glycine 0.1 M and permeabilized in blocking buffer containing 3% BSA, 5% goat serum in 0.2% PBS-Triton X-100. Anti-PABPN1 primary antibody (1/200; ab75855 Abcam) was incubated 1 hour at room temperature in blocking buffer. Cells were washed three times with 0.1% PBS-Triton X-100 prior to incubation with goat anti rabbit secondary antibody conjugated to Alexa Fluor 488 (Life Technologies) together with Phalloidin 555 (Interchim).

### Western blot

Proteins were extracted by sonicating the cells in RIPA buffer (0.15 M NaCl, 0.1% SDS, 50 mM Tris-HCl (pH 8), 2 mM EDTA and 10% Triton-X-100 with protease inhibitor cocktail (Complete, Roche Diagnostics) and phosphatase inhibitor cocktail (Santa Cruz; sc-45064)). Protein concentration was determined by colorimetric detection method (Pierce BCA Protein Assay; Thermo Fisher). Proteins were separated on 4–12% Bis-Tris gels (Invitrogen) and transferred onto a PVDF membrane for 1 h at constant 250 mA at 4°C. Membranes were blocked by incubation in 5% fat free milk or BSA in 1X TBS, 0.1% Tween-20 (TBST) for 1h at room temperature under agitation. Membranes were stained with primary antibodies (PABPN1 Abcam ab75855; eiF2α-P Cell signaling #9721; eiF2α total Cell signaling #9722) overnight at 4°C under agitation. The next day, membranes were washed in TBST and incubated with appropriate secondary antibodies. The G:Box system (Syngene) was used to detect the signals from the membranes.

### RT-PCR and Real-time qRT-PCR

Total RNA was extracted from cell and skeletal muscle samples using Trizol reagent (Invitrogen) according to the manufacturer’s instructions. RNA quality and purity was determined using a ND-1000 NanoDrop spectrophotometer (NanoDrop Technologies). RNA (100 ng for human muscle biopsies, 1 μg for cell pellets and mouse muscle biopsies) was reverse transcribed using M-MLV reverse transcriptase (Invitrogen) according to the manufacturer’s instructions. The splicing of *Xbp1* mRNA was monitored by PCR with 0.5-1 μl of cDNA using Reddy mix polymerase (Thermo Scientific) according to the manufacturer’s instructions. Primers sparing the 26 bp intron sequences are listed in Table S1. The reaction mixture was heated to 94°C for 5 min and followed by 35 PCR cycles: 30 s at 94°C, 30 s at 55°C and 30 s at 72°C followed by a last elongation step at 72°C for 7 min. PCR products from unspliced and alternatively spliced *Xbp1* mRNAs, which are 172 and 146 bp, respectively, were resolved on 5% non-denaturing polyacrylamide or 2% agarose ethydium bromide stained gels. cDNA was used for quantitative PCR reaction using SYBR green mix buffer (LightCycler 480 Sybr green I Master) in a total reaction volume of 9 μl. The PCR reaction was carried out as follows: 8 min at 95°C followed by 50 cycles of the following: 15 s at 95°C, 15 s at 60°C and 15 s at 72°C. Specificity of the PCR products was checked by melting curve analysis using the following program: 65°C increasing by 0. 11°C/s to 97°C. The expression level of each mRNA was normalized to that of human *B2M* mRNA (beta-2 microglobulin) or murine *Rpl0* mRNA (large ribosomal protein, subunit P0) expression. Expression levels were calculated according to the ΔΔCt method. The sequences of primers used for RT-PCR and for real-time qRT-PCR are listed in Table S1.

### Mice

A17 transgenic mice have been previously described (25, 49). Male A17 and wild-type FvB (WT) controls were generated by crossing the heterozygous carrier strain A17 with FvB mice. The mice were genotyped by PCR about four weeks after birth. Animals were housed with food and water ad libitum in minimal disease facilities (Royal Holloway, University of London). Ethical and operational permission for *in vivo* experiments was granted by the RHUL Animal Welfare Committee and the UK Home Office, and this work was conducted under statutory Home Office regulatory, ethics, and licensing procedures, under the Animals (Scientific Procedures) Act 1986 (Project Licence 70/8271).

### Drug administration

For the systemic treatment of OPMD a total of 12 mg/kg/week of GA was administered using either food chow or intraperitoneus injections. The total amount of GA administered per week is in the range of the dose of the GA based Wytensin used in human (tested in clinical trial from ~0.23 to 3.7 mg/kg/week; (73)) for the dose conversion human/mouse see FDA guidance for industry, 2005, Center for Drug Evaluation and Research (CDER), July 2005. For the oral feeding experiment, a 2-week long pilot experiment in FvB healthy mice was conducted to measure the amount of food needed for 13 weeks. The calculation resulted on mice eating an average of 2 pellets (2.8 grams each) per day. For the experiment in OPMD mice, 10-12 week old mice per group (mice fed with GA or bare food) were caged individually and bare or the GA-supplemented pellets (GA was 10mg/kg correspondent to 0.028mg GA/pellet) changed twice a week for 13 weeks until the end of the experiment. During the experiment, food was weighted each time in order to assess the amount of food eaten by the mice. For the intraperitoneal injection, GA was bought from Sigma and diluted in 5% DMSO in saline. Groups of 10-12 week old A17 or FvB mice were injected with either GA 4 mg/kg/injection or with 5% DMSO in saline three times per week for 13 weeks.

### Assessment of functional readouts in living mice

#### Mouse open-field behavioural activity

Soon after the first i.p. injection of GA or saline, activity of mice was evaluated for 60 minutes (12 bins of 5 minutes each) using open field behavioral monitoring and data obtained per each mouse were averaged. Activity cages from Linton Instruments were used and data were collected using Amonlite software (vs 1.4). During the last 2 weeks of treatment with either oral feeding or intraperitoneal injections, the same cages were used to monitor the activity of treated mice according to a protocol designed following the TREAT-NMD guidelines (TREAT-NMD SOP: DMD_M.2.1.002). Briefly, mice were acclimatized to the test chamber for 60 min every day for 4 consecutive days. On the days of data collection, mice were acclimatized for 30 min before data acquisition and data were then acquired every 5 minutes for 1h. This protocol was repeated for 4 days and data obtained per each mouse were averaged. As described in Results, only the first 30 minutes of analyses were considered indicative of the locomotor behavior.

### Grip strength test

The forelimb strength of mice treated with either oral feeding or i.p. injections was assessed using a commercial grip strength monitor (Linton Instrumentation, Norfolk, UK). The protocol described in the TREAT-NMD guidelines was used for the oral feeding experiment (5 tests per mouse over a 3-day period, TREAT-NMD SOP: DMD_M.2.2.001). Mice, held from the tail, were allowed to grab a metal mesh attached to a force transducer and the force produced after a gentle pull, was recorded. 30 seconds were left between each of 5 sequential tests per mouse per day. Data were expressed as gram force per gram of the initial body weight. While behavior of the mice orally fed with drugs was normal and similar along the 3-days long procedure, we observed that all mice treated with chronic i.p. injections (and then manipulated thrice/week by the operator) voluntarily did not grab the mesh and preferred to be dragged during both the second and the third days of analyses, so only the measurements obtained from the 1^st^ day of analysis were used for the study.

### In situ muscle physiology

TA muscle strength was measured by *in situ* muscle physiology performed under terminal anesthesia using a protocol already reported in (74). Briefly 24h after the last injection of saline or GA, muscle function was assessed using tibialis anterior muscles of each mouse. Mice were deeply anesthetized and were carefully monitored throughout the experiment. The distal tendon of the TA muscle was dissected and tied with 4.0 braided surgical silk (Interfocus, Cambridge, UK). The sciatic nerve was exposed and superfluous branches axotomized. The TA tendon was fixed to the lever arm of a 305B dual-mode servomotor transducer (Aurora Scientific, Aurora, Ontario, Canada) via a custom made steel s-hook. Bipolar platinum electrodes located in the distal part of common peroneal nerve, were used to stimulate using supramaximal square-wave pulses of 0.02 ms (701A stimulator; Aurora Scientific) TA muscles. Data acquisition and control of the servomotors were conducted using a Lab-View-based DMC program (Dynamic muscle control and Data Acquisition; Aurora Scientific). Maximum isometric tetanic force (Po) was determined from the plateau of the force–frequency relationship following a series of stimulations at 10, 30, 40, 50, 80, 100, 120, 150 and 180 Hz. Muscle length was measured using digital calipers based on well-defined anatomical landmarks near the knee and the ankle. The specific force (N/cm2) was calculated by dividing Po by TA muscle cross-sectional area. Overall cross sectional area was estimated using the following formula: muscle weight (g)/[TA fiber length (Lf; cm) × 0.6 × 1.06 (g/cm^3^)].

### Samples harvesting, processing and storing

Animals were all culled at the end of the study and after the functional analyses. Tibialis anterior (TA), extensor digitorum longus (EDL), soleus (SOL) and gastrocnemius (GAST) muscles were harvested from mice treated with i.p injections and TA, EDL and SOL muscles were collected from mice treated with oral feeding. Muscles from a leg were snap-frozen in liquid nitrogen for WB and RNA studies and the muscles of the contralateral leg were mounted in corks using OCT and frozen in pre-chilled isopentane in liquid nitrogen for histological analyses. All samples were stored at −80°C before analysis.

### Immunostaining and histological analyses

A Bright OTF 5000 cryostat (Bright Instruments, Huntingdon, UK) was used to section muscles at 10 μm thickness. Muscle sections, placed on coated glass slides (VWR, Lutterworth, UK) were stored at −80°C prior to use. Immunohistochemical staining was performed using the following antibodies: anti-PABPN1 (rabbit monoclonal, diluted 1:100, Abcam ab75855, ON, 4°C), anti-Laminin (rat monoclonal; diluted 1:800; Sigma-Aldrich, 1 h RT), anti-Collagen VI (rabbit monoclonal; diluted 1:400; Abcam ab182744). Alexa fluor (Molecular Probes) antibodies conjugated to 488 or 594 fluorochromes were used to detect the primary antibodies. A published protocol (18, 25, 49) was used to counterstain the sarcolemma with Laminin for PABPN1 immunostaining. Briefly, after fixing sections in Paraformaldeyde 4% the slides were incubated with KCl buffer (1 M KCl, 30 mM HEPES, 65 mM PIPES, 10 mM EDTA, 2 mM MgCl2, pH 6.9) for 1 h at room temperature. Muscle sections were then blocked with 1% normal goat serum in 0.1 M PBS, 0.1% Triton X100 for 1 h and then incubated overnight at 4°C with both PABPN1 and Laminin primary antibodies diluted in the same buffer. Slides were then washed with 0.1% tween 20 in PBS and incubated with fluorophore-conjugated secondary antibodies. Slides were mounted with mounting medium Vector containing 4’,6-diamidino-2-phenylindole (DAPI, 3 μg/ml; Sigma-Aldrich Ltd.) to detect nuclei. For Collagen VI staining 5% milk in PBST was used to block for 1h at RT the unspecific binding and both the primary and secondary antibodies were incubated for 1h at RT in 2.5% fat free milk in PBST. Five, 20X magnification images were randomly taken from a blinded observer and used to calculate the percentage of aggregates, the percentage of centrally nucleated fibres and the area covered by collagen VI. A standard protocol for Sirius red was used to detect the collagen I and III in TA muscles. Two to three 10X magnification images were randomly taken from a blinded observer and used to calculate the area stained by Sirius red and expressed as % on the total area. NIH imageJ analysis software was used to analyze the largest section of each muscle.

### Image acquisition and analysis

Images were visualized using an Olympus BX60 microscope (Olympus Optical, Hamburg, Germany), digitalized using a CCD camera (Photometrics CoolSNAP fx; Roper Scientific) and analyzed using MetaView image analysis system (Universal Imaging, Downington, PA), MetaMorph imaging system (Roper Scientific, Tucson, AZ, USA) software, and ImageJ 1.44o (http://imagej.nih.gov/ij) for quantification analysis.

### Electron microscopy

TA muscles were dissected, cut into small pieces and immediately fixed in 2% glutaraldehyde, 2% paraformaldehyde, 0.1M phosphate buffer. After abundant washes and 2% OsO_4_ postfixation samples were dehydrated at 4°C in graded acetone including a 1% uranyl acetate in 70° acetone step and were finally embedded in Epon resin. Thin (70nm) sections were stained with uranyl acetate and lead citrate, observed using a Philips CM120 electron microscope (Philips Electronics NV) and photographed with a digital SIS Morada camera.

### Gene profile analysis

Enrichment analysis have been performed on the 1099 differentially expressed genes (DEGs) from quadriceps muscles of 26-weeks old A17 mice previously deposited on GEO (GSE26604) (49). The analysis have been done using overexpressed test (75) of clusterProfiler package (76) and genome wide annotation for Mouse (77) in R. The Benjamini & Hochberg correction method has been applied and results with a False Discovery Rate (FDR) lower than 5% and an absolute logFC higher than 0.6 have been kept.

### Statistical Analysis

All data are mean values ± standard error of the mean. Statistical analyses were performed using the Student t-test and the one-way ANOVA with the Bonferroni post-hoc analysis as indicated. GraphPad Prism (version 4.0b; GraphPad Software, San Diego California, USA) was used for the analyses. A difference was considered to be significant at \**P* < 0.05, **P< 0.01 or ***P < 0.001.

